# Disease and climate effects on individuals jointly drive post-reintroduction population dynamics of an endangered amphibian

**DOI:** 10.1101/332114

**Authors:** Maxwell B. Joseph, Roland A. Knapp

## Abstract

The emergence of novel pathogens often has dramatic negative effects on previously unexposed host populations. Subsequent disease can drive populations and even species to extinction. After establishment in populations, pathogens can continue to affect host dynamics, influencing the success or failure of species recovery efforts. However, quantifying the effect of pathogens on host populations in the wild is challenging because individual hosts and their pathogens are difficult to observe. Here we use long-term mark-recapture data to describe the dynamics of reintroduced populations of an endangered amphibian (*Rana sierrae*) and evaluate the success of these recovery efforts in the presence of a recently-emerged pathogen, the amphibian chytrid fungus *Batrachochytrium dendrobatidis*. We find that high *B. dendrobatidis* infection intensities are associated with increases in detectability, reductions in survival, and more infected adults. We also find evidence for intensity-dependent survival, with heavily infected individuals suffering higher mortality. These results highlight the need in disease ecology for probabilistic approaches that account for uncertainty in infection intensity using imperfect observational data. Such approaches can advance the understanding of disease impacts on host population dynamics, and in the current study will improve the effectiveness of species conservation actions.

## Introduction

Amphibians are one of the most threatened groups of vertebrates (Wake and Vredenburg 2008). Although the drivers of amphibian decline vary taxonomically and spatially, the amphibian chytrid fungus *Batrachochytrium dendrobatidis* (Bd) is a major cause of population declines and species extinctions in montane habitats worldwide (Skerratt et al. 2007, Fisher et al. 2009, Grant et al. 2016). In the face of these declines, species recovery will often require introductions to restore extirpated populations, but little is known about the dynamics of population establishment and persistence of threatened amphibians (Armstrong and Seddon 2008), especially in the presence of disease. Given Bd’s role in eliminating populations, we expect introduction outcomes to be shaped by disease impacts on demographic rates, but measuring such impacts in wild populations is difficult (McCallum and Dobson 1995, Briggs et al. 2010).

The mountain yellow-legged frog is emblematic of global amphibian declines. Although formerly abundant in the relatively protected habitats of California’s Sierra Nevada mountains (USA), the two species that make up this taxon (*Rana muscosa* and *Rana sierrae* (Vredenburg et al. 2007)) have disappeared from over 93% of historical localities, due primarily to the introduction of nonnative fish into fishless habitats (Knapp and Matthews 2000, Knapp 2005) and the emergence of Bd (Vredenburg et al. 2010). In response, both species are listed as “endangered” under the U.S. Endangered Species Act and included on the International Union for Conservation of Nature Red List of Threatened Species (IUCN 2017). *R. sierrae* has shown recent signs of recovery in the best-protected portion of its range, including Yosemite National Park, but due to dispersal limitations the re-establishment of populations may require introductions in addition to natural recovery (Knapp et al. 2016). The effectiveness of such actions remains unclear, and the drivers of post-introduction population dynamics and introduction success or failure are poorly understood.

Introduction outcomes may be described in terms of survival, recruitment, and abundance trajectories. Mark-recapture studies provide a method to estimate these quantities in wild populations while accounting for factors that can complicate inference, including imperfect detection (Jolly 1965, Pradel 1996), multiple classes within the population that may be imperfectly resolved (Lebreton and Cefe 2002, Conn and Cooch 2009), and incompletely observed individual-level traits (Royle 2009). This flexibility is critical for understanding disease impacts in wild populations. Multi-state mark-recapture methods have been applied with great success to understand the population-level impacts of Bd, and differences in demographic rates between infected and uninfected classes (Murray et al. 2009, Pilliod et al. 2010, Sapsford et al. 2015, Hudson et al. 2016).

Assuming that survival depends only on whether an individual is infected, and not considering infection intensity (i.e., load), could be problematic given that Bd-caused frog mortality is often a function of load (Briggs et al. 2010). However, individual-level infection data are often not available. For example, Bd load may not be observed if an individual is not encountered or captured on a survey, or if they are captured but no infection data are collected. This presents a major challenge: infection intensity is hard to measure, but may be critical for understanding population dynamics. Previous efforts to understand disease impacts in wild populations as a function of load at the individual level have randomly imputed missing infection load data with the observed distribution of load values (Spitzen-van der Sluijs et al. 2017). However, random imputation could bias inference if load data are not missing at random, for example, if load-dependent disease affects detection.

In this paper, we use a Bayesian multi-state model with data from decade-long mark-recapture studies in two reintroduced frog populations to understand the influence of disease and climate on introduction outcomes. We find that disease dynamics affect introduction success, with high infection intensities leading to more infected adults, a reduction in adult survival, and adults that are easier to detect. Recruitment and uninfected adult survival are also important for population persistence. Differential detectability of adults related to infection intensity points to broader questions about understanding disease impacts in wild populations, and highlights the importance of understanding hidden infection dynamics at the individual level. These insights will inform ongoing and future efforts aimed at restoring the endangered mountain yellow-legged frog, and provide a means to quantitatively assess why some introduction efforts fail and others succeed.

## Methods

### Field sites and methods

The two study lakes are located in eastern Yosemite National Park (California, USA) at elevations of 2880 and 3200 m. *R. sierrae* populations in both lakes were established via translocation from a single nearby donor population (elevation: 3176 m). The donor population has been Bd-positive for at least two decades (Fellers et al. 2001), and contains one of the largest *R. sierrae* populations in Yosemite (minimum population size during this study: 600-1500 adults). The introduction lakes are located 5.5-7.0 km from the donor population and both contain high quality *R. sierrae* habitat (i.e., fishless, 6-12 m deep, with adjacent meadow and stream habitats) (Knapp 2005). Because these lakes harbor a sensitive species, we refer to them by the pseudonyms Alpine and Subalpine (Lindenmayer and Scheele 2017). Visual encounter surveys (Knapp 2005) conducted just prior to the first introductions indicated that both lakes lacked *R. sierrae* of any life stage. Prior to introduction, adult frogs (*≥* 40 mm snout-vent length (SVL)) were captured at the donor site, and tagged (8mm passive integrated transponder (PIT) tags), measured, and weighed. To estimate Bd load, we also collected a skin swab from each frog using standard methods (Hyatt et al. 2007, Vredenburg et al. 2010).

The initial introduction of frogs to Alpine and Subalpine occurred during summer 2006 and 2008, respectively, and additional introductions to supplement both populations were conducted in subsequent years. In the first introduction to Alpine, frogs were transported on foot; in all subsequent introductions to both Alpine and Subalpine, frogs were transported via helicopter (Table 1).

**Table 1.** Schedule of frog introductions to the study lakes.

| Lake | Year | Number of frogs |
| --- | --- | --- |
| Alpine | 2006 | 40 |
| . | 2013 | 20 |
| Subalpine | 2008 | 36 |
| . | 2013 | 19 |
| . | 2015 | 20 |
| . | 2017 | 30 |

To describe the dynamics of the introduced populations, both populations were assessed for 10-12 years using mark-recapture methods. Between 2006 and 2012, lakes were visited approximately once per month during the summer active season (June-September) and on a single day (primary period) all habitats were searched repeatedly for frogs which were captured using hand-held nets. Adult frogs were identified via their PIT tag (or tagged if they were untagged), measured, weighed, swabbed, and released at the capture location. During 2013-2017, we adopted a robust design in which all habitats were searched during several consecutive days (surveys), and frogs processed as described above. Within a primary period, frogs that were captured on more than one survey were measured, weighed, and swabbed only when first captured.

Skin swabs were analysed using standard Bd DNA extraction and qPCR methods (Boyle et al. 2004) except that swab extracts were analyzed singly instead of in triplicate (Kriger et al. 2006, Vredenburg et al. (2010)). During 2005-2014, we used standards developed from known concentrations of zoospores (Boyle et al. 2004) and after 2014 we used standards based on single ITS1 PCR-amplicons (Longo et al. 2013). Based on paired comparisons between samples analyzed using both types of standards, Bd in the study area has an average of 60 ITS1 copies per zoospore. To express all qPCR results as the number of ITS1 copies, starting quantities obtained using the zoospore standard (measured as “zoospore equivalents”) were multiplied by 60. In addition, all qPCR quantities (regardless of standard) were multiplied by 80 to account for the fact that DNA extracts from swabs were diluted 80-fold during extraction and PCR (Vredenburg et al. 2010).

We acquired hourly air temperature and daily snow depth data from two nearby meteorological stations (ERY and DAN, respectively) and snow water equivalent data from a manually-measured snow course (DAN; California Data Exchange Center - http://cdec.water.ca.gov). The stations and snow course are 5-17 km from the study lakes. We used snow water equivalent as measured on April 1 of each year as a measure of winter severity, to be used as a covariate for frog recruitment and survival. Daily snow depth data were acquired to model survey occurrence as described below. To better understand the detection process, we averaged hourly air temperature data collected between 0900-1600 on each day a survey took place, to derive a “survey air temperature” metric.

### Model development

We developed a hierarchical Bayesian hidden Markov model to understand how environmental factors and partly observed time-varying individual traits (Bd loads) jointly drive population dynamics. This model also needed to account for the partly deterministic recruitment process that arises with introductions. The result is an extended open population Jolly-Seber mark-recapture model with known additions to the population (introduced adults), continuous uncertain disease states of individuals which may or may not be captured, and sampling that is unevenly distributed in time.

In a small number of cases, adults were captured, weighed, and measured, but no swab data were associated with the capture (e.g., if a swab was lost). In these cases (3% of 2697 swabs), capture data indicate that the adult was alive during the survey, but its infection status is unknown. As such, there are three possible observations for captured frogs: detected/infected, detected/uninfected, and detected/unknown infection status. A fourth possible observation class corresponds to non-detection: an individual was not observed during a survey.

### Mark-recapture model structure

We consider four possible observations for each individual *i* = 1, *…, M* on each survey *j* = 1, *…, n*_*j*_: *o*_*i,j*_ = 1 indicates that the individual was not detected; *o*_*i,j*_ = 2 indicates that the individual was observed and the swab collected for that individual indicated that they were uninfected; *o*_*i,j*_ = 3 indicates that the individual was observed and results from the collected swab showed that the frog had a non-zero Bd load; and *o*_*i,j*_ = 4 indicates that the individual was observed, but no swab data were available from the capture.

We use parameter-expanded data augmentation to account for the fact that the total number of adults in the population is unknown (Royle and Dorazio 2012). Across the entire time period of the study, we assume *N*_*s*_ individuals have existed in the population, comprising the superpopulation of individuals that were present at some time. While *N*_*s*_ is not directly observable, across all surveys *n ≤ N*_*s*_ unique individuals were known to be present in the population (because they were introduced and/or captured). An estimate of *N*_*s*_ can be acquired by considering a large number *M > N*_*s*_ of individuals, *M − N*_*s*_ of which never existed (Royle 2009). Thus, the model has a state corresponding to individuals that have not recruited yet. Here, *M* was chosen to be 2218 at Alpine (618 observed unique individuals plus 1600 augmented individuals), and 558 at Subalpine (158 observed individuals, and 400 augmented individuals). These values were chosen to be considerably greater than our prior guess of *N*_*s*_, and we also verified that posterior estimates of *N*_*s*_ were much less than *M* to avoid problems on the boundary of this augmented parameter space (Dennis et al. 2015).

We denote the true state of individual *i* in primary period *t* as *u*_*i,t*_, for every individual *i* = 1, *…, M* and each primary period *t* = 1, *…, n*_*t*_. The four states that we consider are: *u*_*i,t*_ = 1 for individuals that have not recruited, *u*_*i,t*_ = 2 for uninfected adults, *u*_*i,t*_ = 3 for infected adults, and *u*_*i,t*_ = 4 for dead individuals. Each survey *j* = 1, *…, n*_*j*_ occurs in one of the *n*_*t*_ primary periods, and we denote the primary period in which survey *j* takes place as *t*_*j*_. Each primary period *t* occurs within one year, but within a year there are multiple primary periods. We set the year containing the first primary period to *y*_*t*=1_ = 1, and generally *y*_*t*_ represents the year containing primary period *t*. Years increment by one until the final year of the mark recapture efforts, which we denote *n*_*y*_: *y* ∈ {1, 2,…, *n*_*y*_}. We assume that within a primary period, the state of each individual does not change (i.e., individuals do not recruit into the adult population, gain or lose Bd infection, or die). This assumption is justified by the short time intervals between surveys within primary periods.

### Observation model

An emission matrix Ω_*i,j*_ links observations to hidden states, with the elements of Ω_*i,j*_ providing the probability of each possible observation of individual *i* in survey *j*, given the true hidden state. The rows in Ω_*i,j*_ correspond to the state of individual *i* in primary period *t*_*j*_, and the columns correspond to an observation of individual *i* in survey *j*:

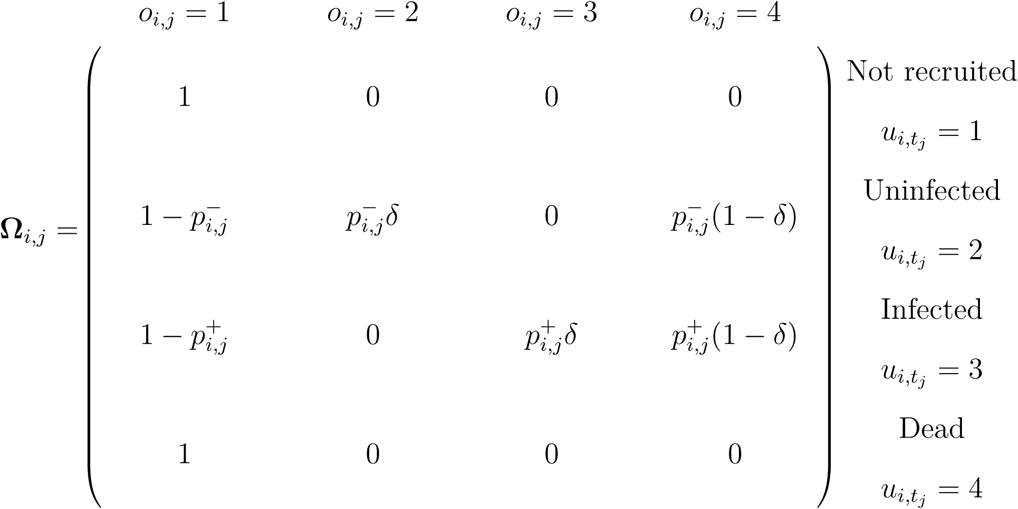

The structure of Ω_*i,j*_ implies that there are no mistaken individual identifications (those that are dead or not recruited are never detected), and that a swab successfully makes it to the lab and provides qPCR data with probability *δ*, conditional on the animal being detected. Detection probabilities are provided by 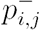 for uninfected (Bd negative) and 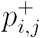 for infected (Bd positive) individuals, and these detection probabilities can vary by individual and survey. We also assume that there are no false-positive or false-negative Bd results (Hyatt et al. 2007), though relaxing this assumption may be a promising future area, given the potential sensitivity of swab results to infection intensity (Miller et al. 2012).

### State model

The hidden states of each individual evolve as a Markov process with transition matrix Ψ_*i,t*_, the entries of which provide the probability of transitioning to state *u*_*i,t*+1_ (the column index) from state *u*_*i,t*_ (the row index). This matrix is different for individuals that naturally recruit versus those that recruit deterministically due to introduction. For naturally recruiting individuals:

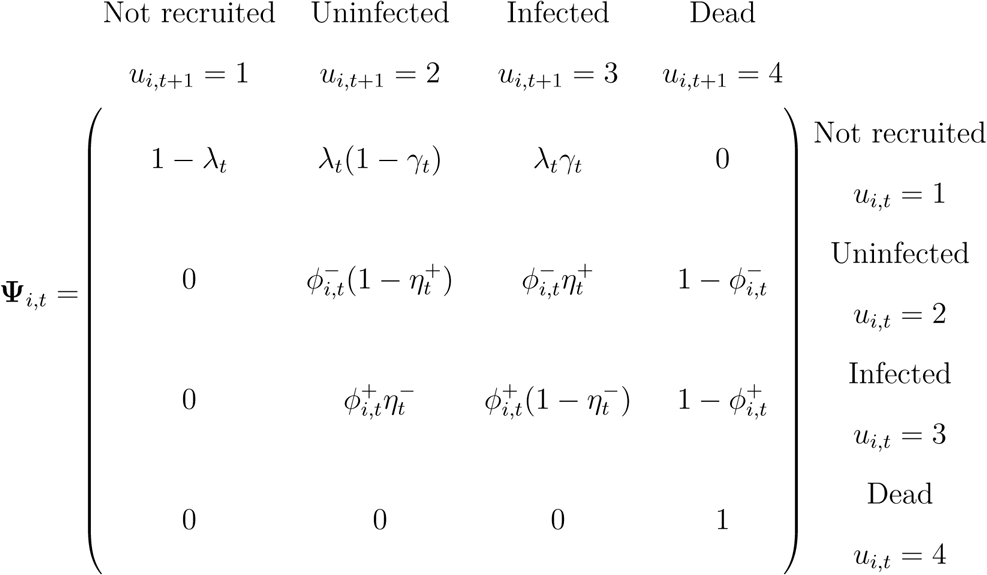

where λ_*t*_ is the probability that an individual enters the adult population between primary periods *t* and *t* + 1, *γ*_*t*_ is the probability that a recruiting individual is infected conditional on entry, 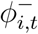 is a survival probability for an uninfected adult, 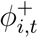 is a survival probability for an infected adult, 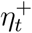 is the probability of transitioning from the uninfected to infected class conditional on survival, and 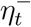 is the probability of transitioning from the infected to uninfected class conditional on survival.

For introduced individuals, the recruitment process is deterministic. Specifically, for each introduced adult, there is zero chance that they recruit prior to the primary period in which they are introduced, and if they are introduced at time *t^intro^_i_*, then the probability that they recruit into a particular class (their state upon introduction) must be one. For these introductions, all introduced adults were infected, and thus recruited into the infected adult class, leading to the following transition matrix for introduced animals, where the recruitment process is completely determined by *t*^intro^_*i*_, such that Ψ_*i,t*_ is:

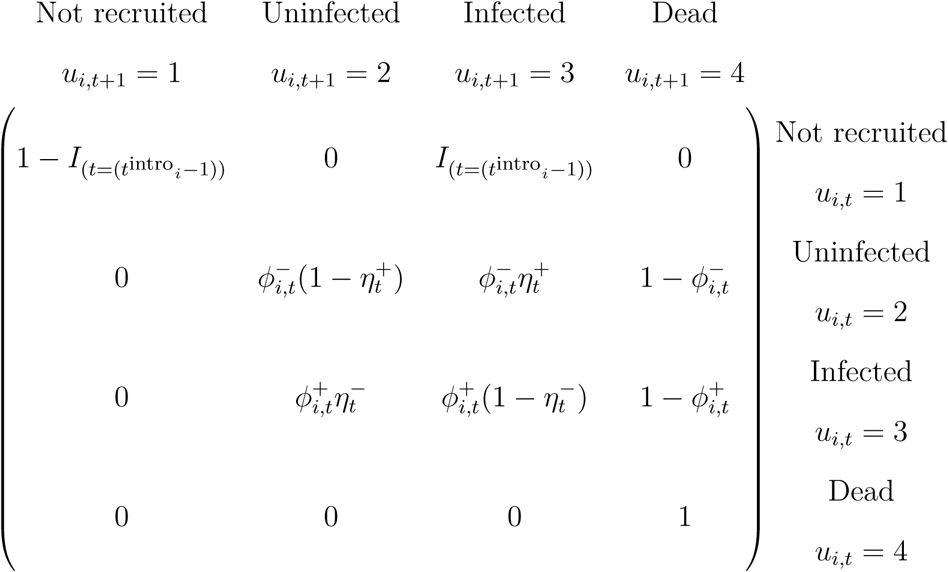

where *I*_(*t*=(*t*^intro^*i-*1))_ is an indicator function equal to one when *t* is equal to *t^intro^_i_*− 1. An imaginary first primary period augments the collection of primary periods when introductions took place and when surveys occurred, and was set to occur one week before the initial introductions into each population. For this augmented period *t* = 1, we assume that all individuals are in the “not-recruited” class, *u*_*i,*1_ = 1 for *i* = 1, *…, M* (Figure 1). Given that neither lake was known to contain adult frogs immediately prior to introduction, this is potentially a fair assumption, but in case the assumption was violated and adults were present prior to the introduction, we include a time-varying adjustment into the recruitment model (described below).

**Figure 1.**
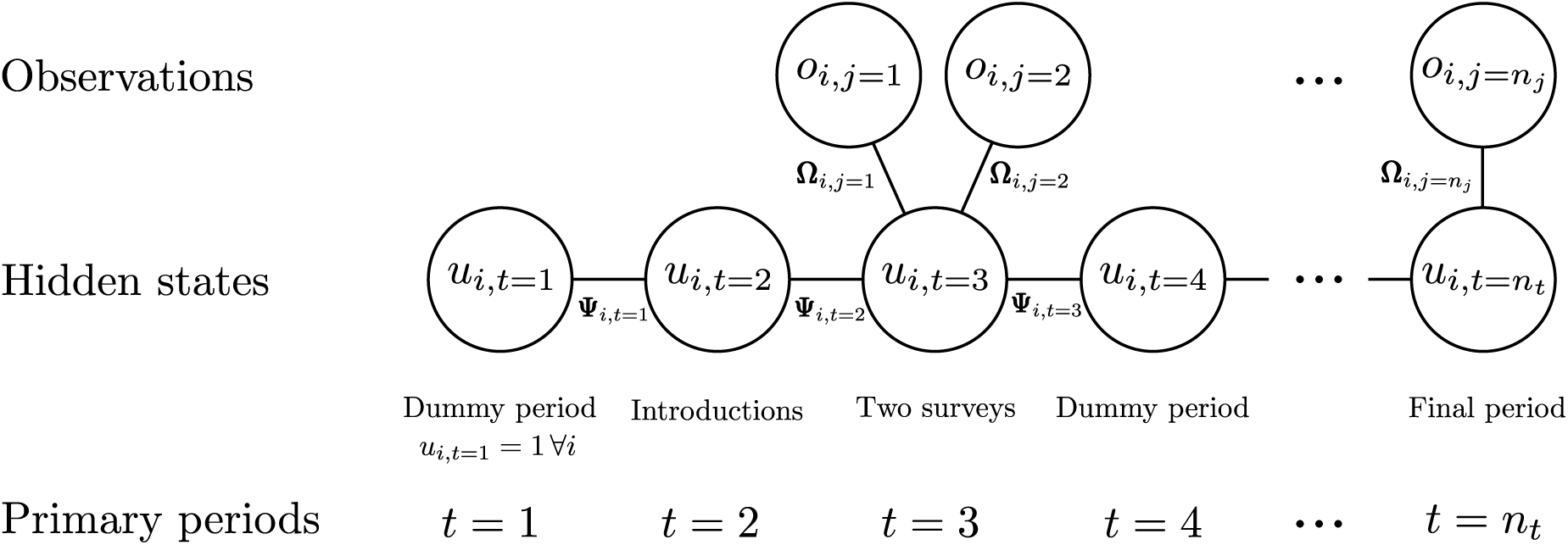
Summary of hidden Markov model structure. On the first primary period *t* = 1, we assume that all individuals *i* = 1, *…, M* are in the not-recruited class. The second primary period represents the first introduction. In this example, two surveys *j* = 1, 2 are conducted on the third primary period. The fourth primary period is a dummy period, inserted to increase the regularity of time intervals between primary periods, and has no associated surveys.

The number of primary periods was not uniform across survey years, and as such, the time between primary periods was also heterogeneous, complicating the interpretation of transition probabilities. Thus, at both sites, we used augmented primary periods to increase the regularity of time intervals between primary periods. These were chosen to occur when primary periods might have occurred, e.g., not during the winter months when the lakes are snow-covered, using a statistical model of when surveys are conducted. At each site, we fit a binomial generalized additive model with thin plate spline smoothers for day-of-year and daily snow depth at Dana Meadows, where the response variable was zero or one for each day, with one indicating that a survey took place. These models were fit using the gam function in the mgcv package in the R programming language (R Core Team 2017, Wood 2017). Then, we predicted new values from these models, with the goal of augmenting the set of primary periods for each year at each site until all years except the first and last had the same number of primary periods (the maximum number of primary periods within one year at each site: ten at Alpine and eight at Subalpine). Augmented primary periods were only accepted if there was no primary period within five days (before and after) of the proposed augmented primary period. The end result is a set of primary periods that are more uniformly distributed in time than the original collection of empirical primary periods, though these augmented primary periods are not associated with real surveys.

### Parameter model

#### Detection probabilities

Among uninfected adult frogs, we assume that the probability of detection varies with survey air temperature (Sinsch 1984), so that

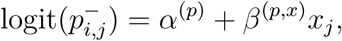

where *α*^(*p*)^ is an intercept parameter, *β*^(*p,x*)^ is the effect of survey air temperature on detection probability, and *x*_*j*_ is the survey air temperature for survey *j*, for all *i, j*. Among infected adult frogs, the detection model was expanded to allow for an adjustment to account for being infected, and a further adjustment to deal with variation due to Bd load:

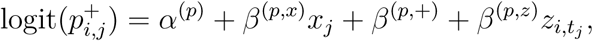

where *β*^(*p,*+)^ is the adjustment on the intercept for infected adults, and *β*^(*p,z*)^ is a coefficient for the log Bd load 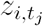 of individual *i* in primary period *t* containing survey *j*, for all *i, j*.

#### Recruitment probabilities

The recruitment model was designed to account for annual variation in Bd loads, whether primary periods spanned years, and winter severity. For the probability of entering the population between primary period *t* and *t* + 1, we have:

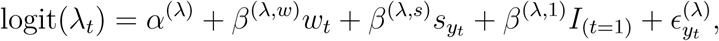

where *α*^(λ)^ is an intercept term, and the effect of an overwinter transition is represented as *β*^(*λ,w*)^, with *w*_*t*_ as a binary indicator of whether a transition from period *t* to *t* + 1 spans a winter (or equivalently, two years). The effect of the previous winter’s severity is *β*^(λ,*s*)^, where *s*_*y*_ is previous winter’s severity. The parameter *β*^(λ,1)^ is an adjustment for the recruitment probability after the first imaginary primary period, which could account for undetected individuals present prior to introduction. Finally, 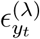 is an adjustment to account for extra annual variation. The probability that an individual is infected, conditional on recruitment is modeled as a function of expected Bd load among infected adults:

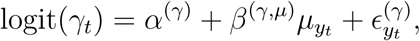

where *α*^(*γ*)^ is an intercept, *β*^(*γ,µ*)^ represents the effect of mean Bd load among infected adults in the year containing primary period *t* (denoted *µ*_*y*_), and *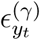* is a year-specific adjustment.

#### Survival probabilities

Survival of uninfected adults was modeled as a function of whether a transition spanned an overwinter period and winter severity:

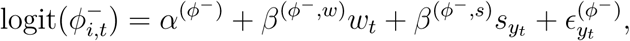

where *α*^(*φ*−)^ is an intercept parameter, *β*^(*ø*−^*,w*) is an adjustment for overwinter transitions, *β*^(*φ*−^*,s*) is a coefficient for winter severity, and 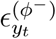 is a year-specific adjustment. The survival probabilities for infected adults were similarly modeled, but with additional effects of individual Bd load:

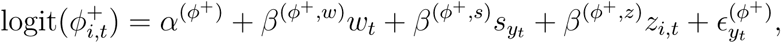

where *z*_*i,t*_ is the log transformed Bd load of individual *i* during primary period *t, β*^(*ø*+^,*z*) is a coefficient for Bd load, and the remainder of parameters are defined using the same notation as for the survival of uninfected adults.

#### Loss and gain of infection probabilities

The probability that an infected adult loses infection was modeled as a function of mean Bd load in the infected population, and whether a transition occurred from one year to the next:

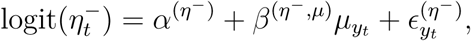

where *α*(*η*^-^) is an intercept, *β*(*η*^-^,µ) is the effect of expected Bd load among infected adults, and *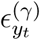* is a year-specific adjustment. Transitions from the uninfected to infected class were modeled similarly:

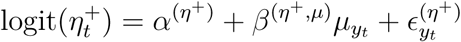

where parameter notation conventions and definitions match those for transitions from the infected to uninfected classes.

#### Bd loads

The fact that individuals are imperfectly detected, and that occasionally, individuals are detected but no disease information is recorded presents a challenge for including individual-level Bd loads in the model. Within the infected population, Bd loads are partially observed when individuals are captured and swabs are collected. We used a normal distribution to represent (potential) log Bd loads:

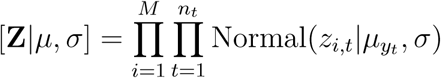

When the load of individual *i* during primary period *t* was observed, this specifies a likelihood. Otherwise, this specifies a prior distribution for the potential log load of individual *i* on primary period *t*, conditional on infection. The expected value of Bd load among infected adults was assumed to vary among years, and potentially vary as a function of winter severity:

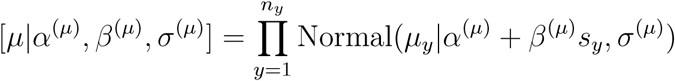

#### Prior distributions

Based on knowledge of the observation process, we expected that most captures resulted in a non-missing swab result, leading to the specification for the prior on δ as [δ] = Beta(*δ|*9, 1). Annual adjustments were modeled using zero-mean normal distributions with unknown standard deviations specific to the process of interest, e.g., for the probability of entering the population: 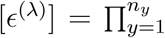 Normal 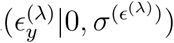. Standard deviation parameters were given half normal priors i.e., 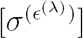 Normal 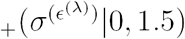, and all remaining parameters were given Normal(0, 1.5) priors. The full factorization of the joint distribution of data and parameters is provided in Appendix A: Joint distribution factorization.

#### Parameter estimation

We implemented the model in Stan, a probabilistic programming language, and sampled from the approximate posterior distribution of parameters with automatic differentiation variational inference (Kucukelbir et al. 2015, Carpenter et al. 2016). We used the forward algorithm (Zucchini et al. 2016) to circumvent the need to sample from the discrete space of true individual states (see Appendix B: Forward algorithm description for details). Variational inference uses a simple family of distributions as an approximation of the posterior distribution, optimizing parameters of this simpler approximation (the variational distribution) to minimize the Kullback-Leibler divergence between the true posterior and the variational distribution. A variational approximation was necessary for the models to run in a reasonable amount of time (≈ 2 - 4 hours) on finite resources (r4.2xlarge EC2 instances on Amazon Web Services, with 8 virtual CPUs and 61 GiB RAM). To verify our model implementation, we simulated data with known parameters from the model to evaluate parameter recovery, and check that parameters were identifiable. These simulations allowed us to identify a lack of identifiability in an earlier model specification which included multiplicative interaction terms between winter severity and Bd load for both the probability of entering the population, and the probability that new adults were infected conditional on recruitment.

All code and data necessary to reproduce the analysis and manuscript is publicly available at https://www.github.com/snarl1/sierra-reintroduction-cmr (Joseph and Knapp 2018). The workflow is wrapped into GNU Make command (Stallman et al. 2004), the manuscript is written in R Markdown (Allaire et al. 2018), and we used the R programming language (R Core Team 2017) with the assertthat, ggrepel, ggridges, ggthemes, lubridate, mgcv, patchwork, reshape2, rstan, and tidyverse packages to facilitate data processing, model fitting, and visualization (Wood 2004, Wickham 2007, 2017a, 2017b, Grolemund and Wickham 2011, Stan Development Team 2016, Pedersen 2017, Slowikowski 2017, Arnold 2018, Wilke 2018). The computational environment with these dependencies is containerized via Docker, and the Dockerfile for the image exists in the GitHub repository for this project (Boettiger 2015).

## Results

At both sites, infected adults were easier to detect than uninfected adults, and detection probabilities also increased with survey air temperature (Figure 2A). For a survey with an air temperature of 17 °C, the estimated probability of detecting an uninfected adult at Alpine was 0.157 (0.15, 0.163; posterior median and 90% CI), but for an infected adult with average Bd load, the probability of detection was 0.324 (0.306, 0.34). At Subalpine, for a survey with the same air temperature, the probability of detecting an uninfected adult was 0.19 (0.17, 0.211), but the probability of detecting an infected adult with an average Bd load was 0.598 (0.551, 0.646). Among infected adults, there was evidence for additional increases in detectability with increases in Bd load in the Alpine population (posterior median for *β*^(*p,z*)^: 0.107, 90% CI: 0.086, 0.129) but not in the Subalpine population (*β*^(*p,z*)^: 0.001, 90% CI: −0.056, 0.053) (Figure 2B).

**Figure 2.**
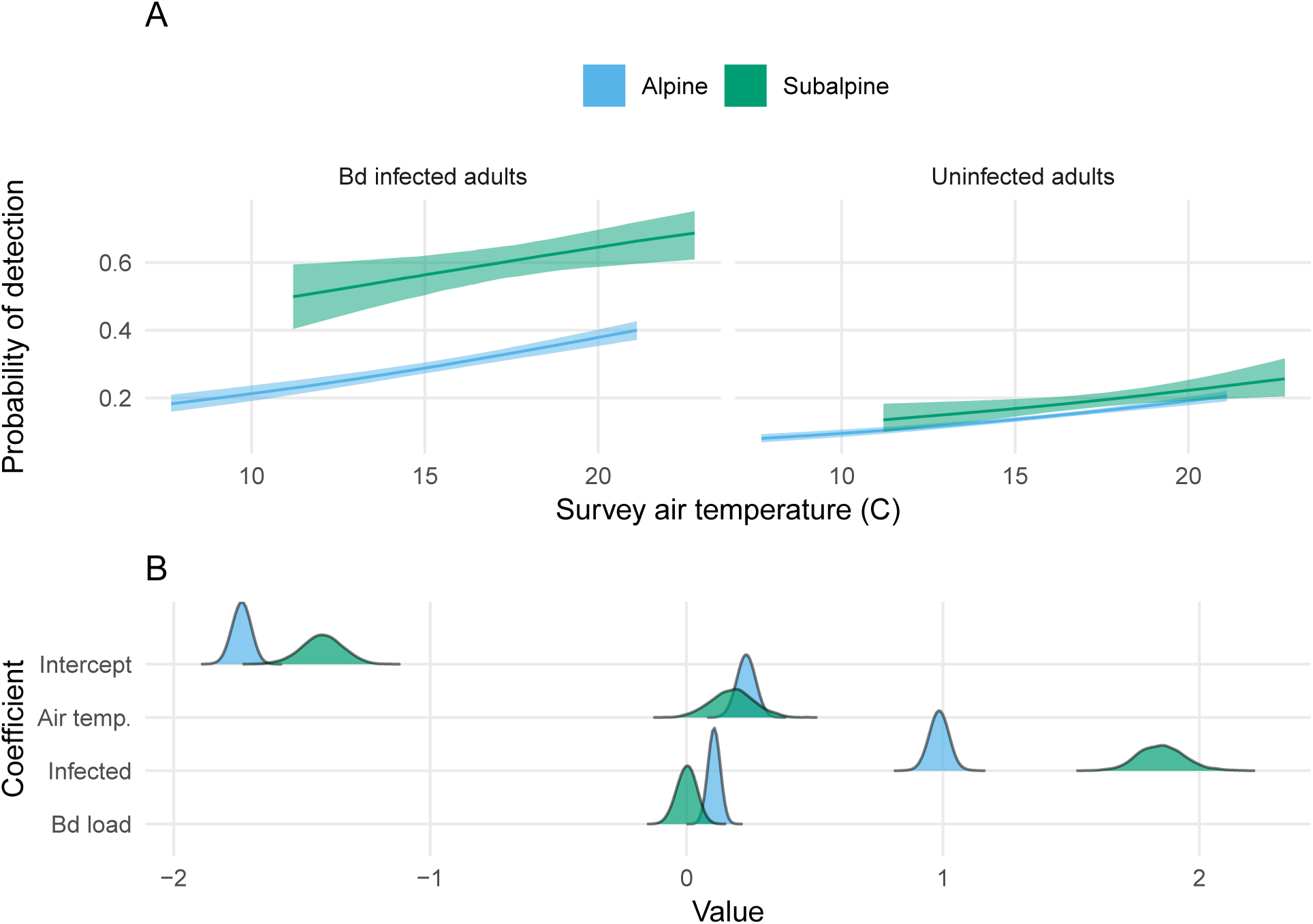
A. Estimated detection probabilities as a function of survey air temperature for Bd-infected and uninfected adults. The ribbons represent the 90% posterior credible interval, and the line represents the posterior median. Predictions for Bd infected adults correspond to individuals with average infection loads at each site. B. Parameter estimates for the detection model. Each of the detection parameters are shown on the y-axis. Color indicates population.

Despite the two study populations being 12 km apart, mean Bd loads among infected adults varied synchronously between populations. After initial introduction in 2006 or 2008, mean loads at both sites were typically between 1,000 and 10,000 copies, with a reduction in mean Bd load during the year following the first introduction (Figure 3). Thereafter, loads were relatively low and stable through 2012. At both sites mean loads were uncharacteristically low in 2013 but increased in subsequent years until reaching a peak in 2016. Mean Bd loads at Subalpine tended to be higher than at Alpine, particularly in the period from 2015 − 2017. The estimated correlation over time between mean log Bd loads at the two study sites was 0.636 (0.315, 0.861). This correlation was not due to a shared response to winter severity because winter severity did not influence Bd load at either site (at Alpine *β*^(*µ*)^: −0.12 (−0.604, 0.37), at Subalpine *β*^(*µ*)^: −0.022 (−0.909, 0.912).

**Figure 3.**
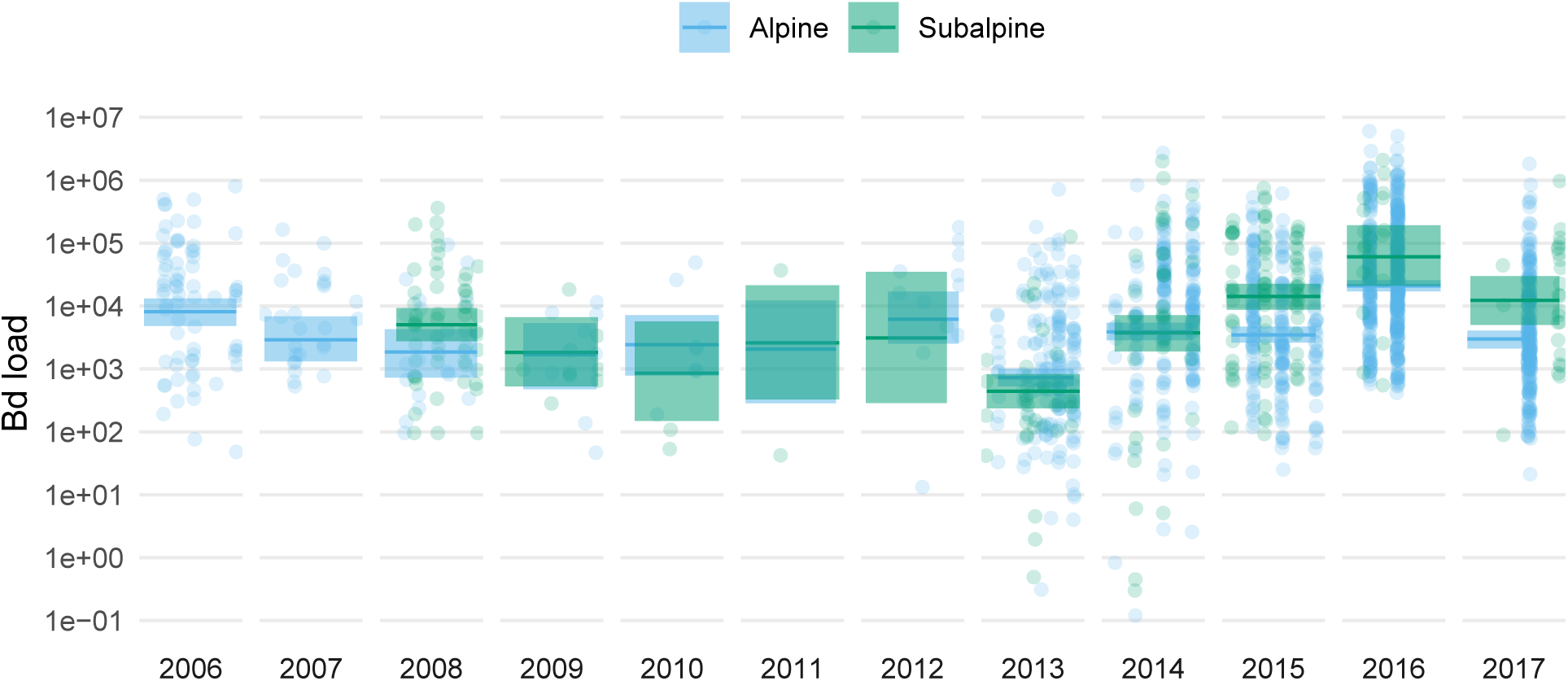
*Batrachochytrium dendrobatidis* (Bd) infection loads over time. Observed load values from skin swabs are shown as points on a log10 scale, colored by site. The estimated average Bd infection load is shown as a line (posterior median) with a 90% credible interval ribbon.

Following initial introduction, both sites experienced multiple years of low abundance, but in 2013 the adult population at Alpine began increasing, reaching values upwards of 400 adults by late-summer 2016 (Figure 4A). In contrast, at Subalpine, the adult population probably has not exceeded 50 individuals over much of the study duration, and introduction events account for the largest recruitment pulses. At both sites, infected adults tended to outnumber uninfected adults throughout the study period. The total number of adults to have ever existed (*N*_*s*_) was estimated to be 768 (686, 848) at Alpine, and 172 (147, 212) at Subalpine.

**Figure 4.**
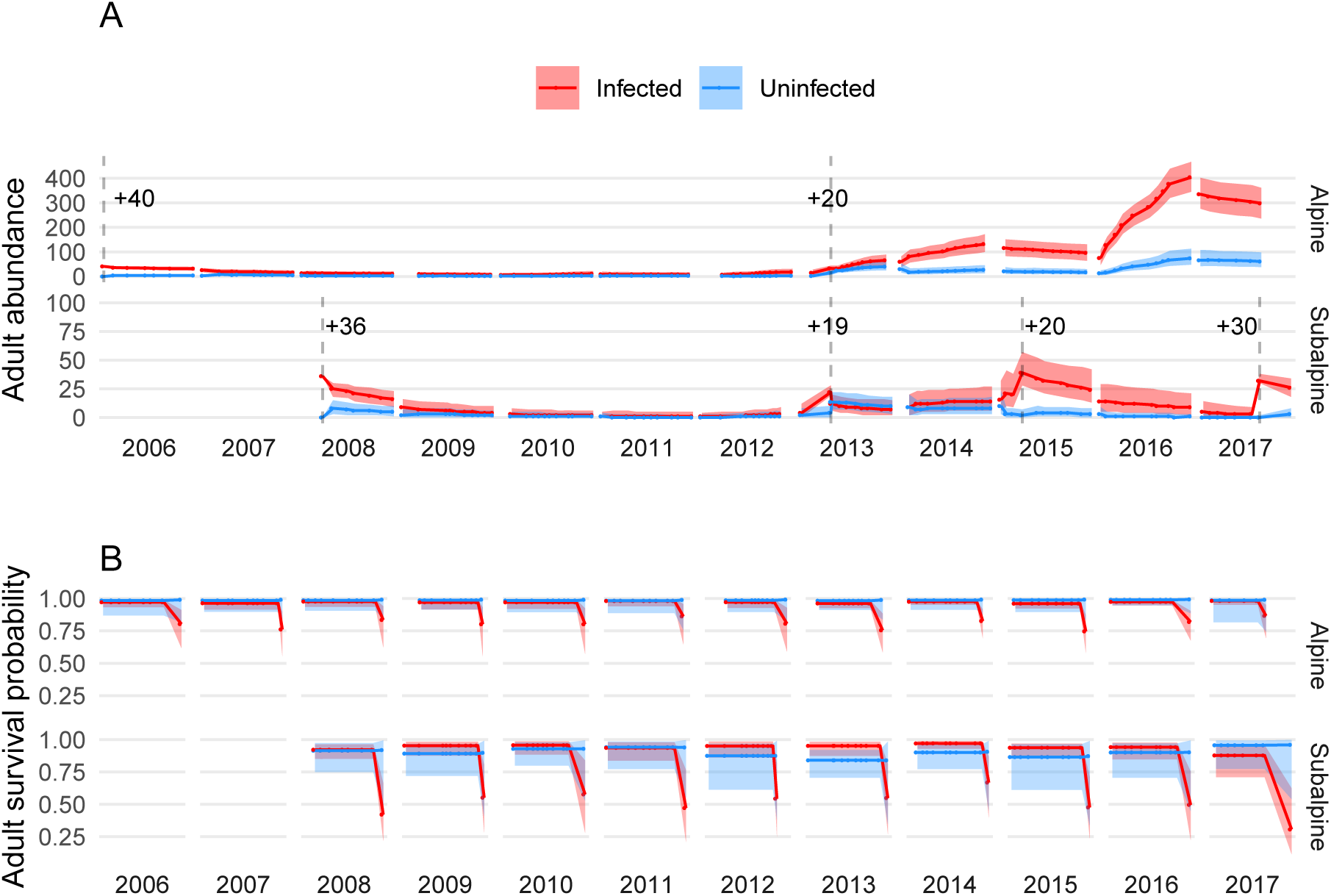
Estimated abundance (A) and survival (B) time series for infected and uninfected adults at both populations. Time is shown on the x-axis, with facets for years. Posterior medians for each primary period are shown as points connected by line segments, and the 90% credible interval is shown as a shaded ribbon. In A, the vertical dashed lines indicate introduction events, and the number of adults added in each event is indicated with a “+” symbol. Also note the difference in the y-axis scales between the two sites.

Infected adults experienced decreased overwinter survival, especially in the Subalpine population, unlike uninfected adults which maintained overwinter survival that was comparable to within-summer survival (Figure 4B). Bd load reduced infected adult survival at Alpine (Figure 5), with *β*^(*ø*+^,*z*) estimated to be −0.132 (−0.211, −0.045) at Alpine, and −0.078 (−0.199, 0.05) at Subalpine. Winter severity appeared to increase survival of Bd infected adults at Alpine (*β*^(*ø*+^,*s*): 0.303 (0.173, 0.436), but may have been associated with lower survival at Subalpine (−0.239 (−0.465, 0)). At both sites, winter severity coefficients overlapped zero for uninfected adults (*β*^(*ø*−^*,s*): −0.044 (−0.735, 0.661) at Alpine, and 0.349 (−0.139, 0.842) at Subalpine). Overall, uninfected adults had higher survival at Alpine (*α*^(*φ−*)^: 4.146 (3.42, 4.883)) than Subalpine (*α*^(*ø−*)^: 2.183 (1.782, 2.594)) (Figure 4B).

**Figure 5.**
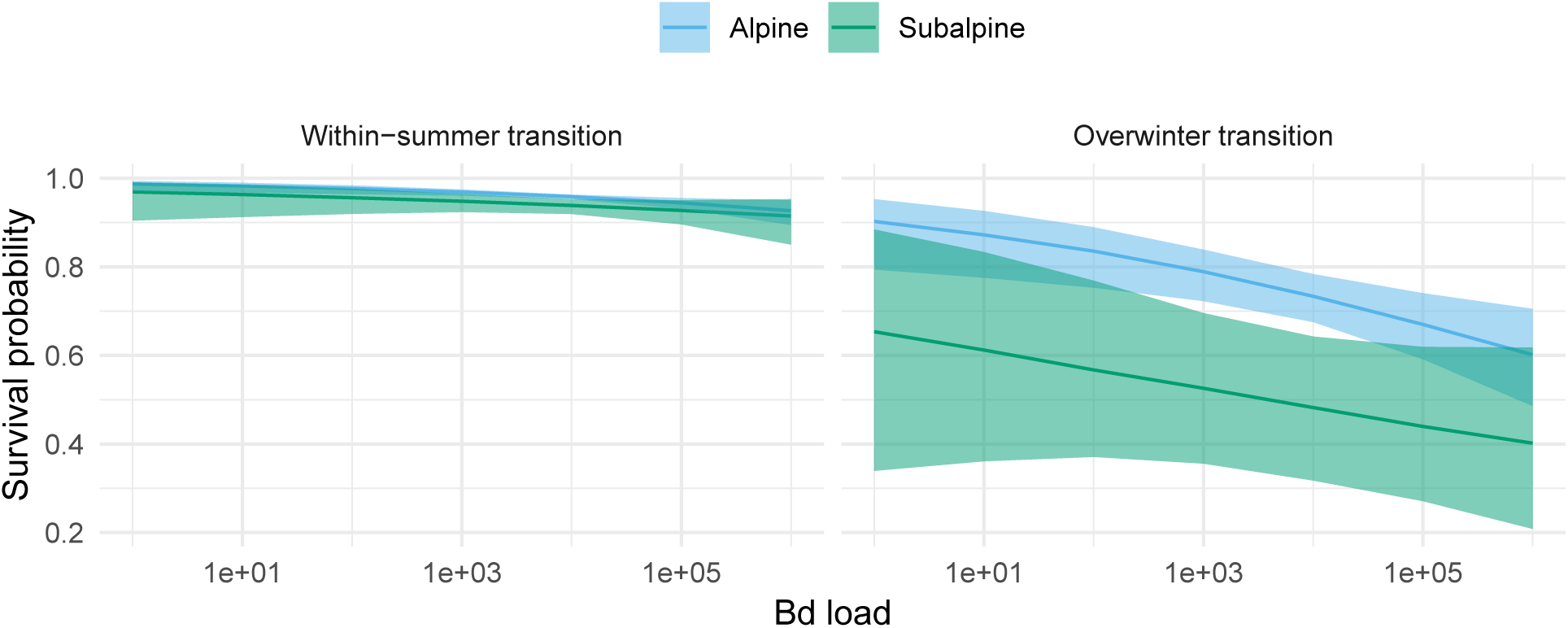
Estimated survival probabilities for within-summer and overwinter transitions among infected adults as a function of Bd infection load. The x-axis shows Bd load over the range of observed values. The y-axis represents survival probabilities. Posterior medians are lines and 90% credible intervals are ribbons.

Introduced adults survived longer on average at Alpine compared to Subalpine (Figure 6). At the end of the summer following the initial introduction of adults at Alpine, the proportion of adults surviving was 0.825 (0.675, 0.95). At Subalpine, only about half (0.583 (0.385, 0.778)) of adults survived to the end of the first summer following initial introduction. In the year after the initial introduction, the estimated proportion of introduced adults surviving at Alpine was 0.675 (0.45, 0.85), and at Subalpine just 0.306 (0.139, 0.528). Similar differences in survival were evident following subsequent introductions, with low within-summer and among-year survival at Subalpine, and a higher survival rate at Alpine (Figure 6).

**Figure 6.**
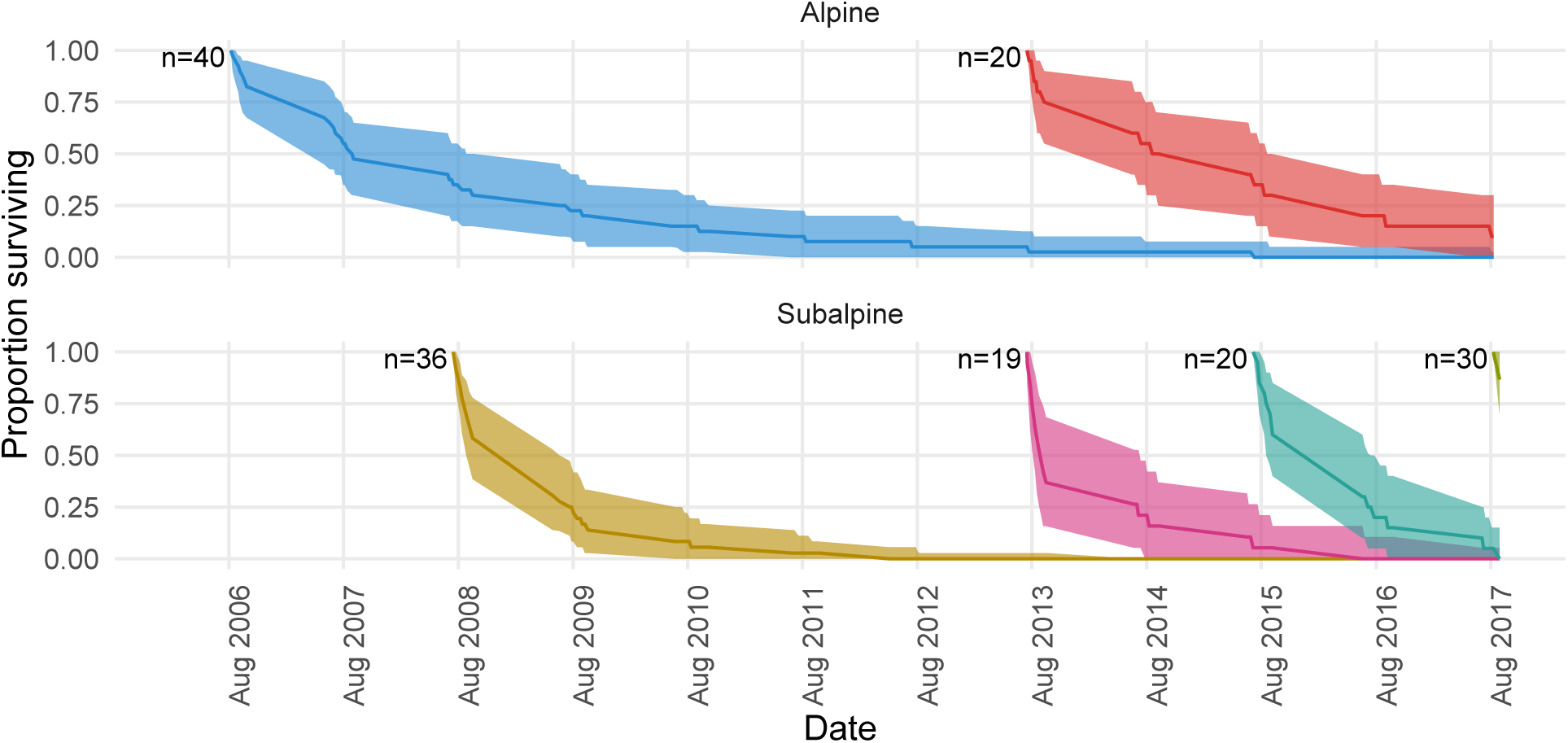
Estimated survival of introduced adults at both populations, colored by introduction event. Time is shown on the x-axis, with the proportion of adults surviving on the y-axis. The number of individuals introduced in each introduction event is labeled as (n=…). The ribbons represent the 90% posterior credible interval, with the midline representing the posterior median.

Little to no recruitment was observed during the 3-4 years following the initial introductions, consistent with the multi-year larval and juvenile stages in this species (“recruitment” is the addition of new adults to the population). However, Alpine experienced large recruitment pulses in 2013, 2014, and 2016 (Figure 7). The Subalpine population had smaller pulses of recruitment during the period 4-6 years after the introduction, but little recruitment in subsequent years (Figure 7). Recruitment pulses were asynchronous between the two populations, particularly in 2016 when a very large pulse of recruits was observed at Alpine but not Subalpine.

**Figure 7.**
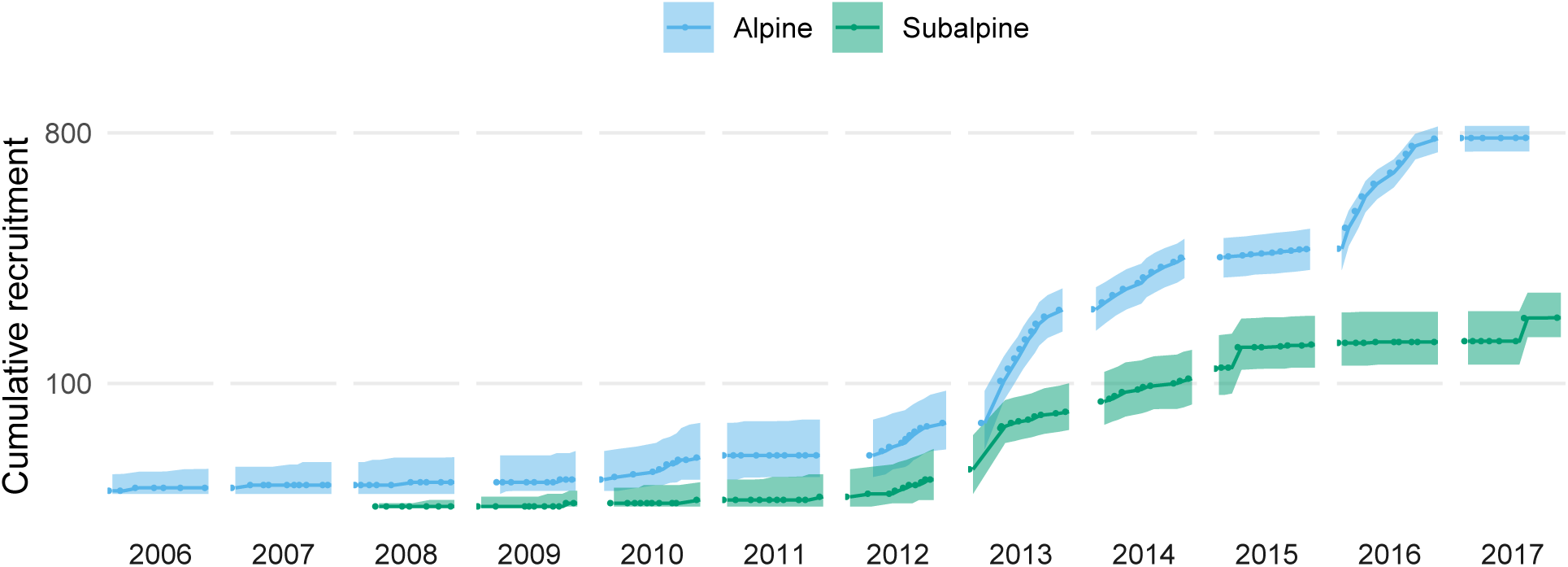
Estimated cumulative number of natural recruits into the adult population at both sites over time (on a log10 scale). The ribbons represent the 90% posterior credible interval, with the midline representing the posterior median. Dots connected by lines represent primary periods.

Recruitment dynamics varied as a function of winter severity and Bd load. In both populations, winter severity reduced recruitment (at Alpine *β*^(λ,*s*)^: −1.41 (−1.55, −1.264), at Subalpine *β*^(λ,*s*)^: −1.295 (−1.612, −1.001)). Recruitment overwinter (before the first primary period of the year) was less likely than recruitment among within-year primary periods at Alpine (*β*^(λ,*w*)^: −2.763 (−3.946, −1.551)), but more likely at Subalpine (*β*^(λ,*w*)^: 1.677 (1.117, 2.256)). Conditional on entering the population, adults at Alpine were more likely to recruit into the infected class when mean Bd loads were high (*β*^(*γ,µ*)^: 1.195 (0.356, 2.104)), but mean Bd load was not associated with the probability of recruiting as infected at Subalpine (*β*^(*γ,µ*)^: −0.575 (−2.199, 1.192)), perhaps due to low recruitment and therefore low power to detect such an effect.

At both sites, transitioning from the uninfected to the infected adult class (gaining infection) was more likely than transitioning from the infected to uninfected class (losing infection) over most of the study period. Among-year variation in transition probabilities was similar at both sites, with 2013 having higher than average loss-of-infection probabilities (Figure 8A). This was most likely due to exceptionally low mean Bd loads in 2013: when Bd loads were low, adults were more likely to lose infection at both study sites, and less likely to gain infection in the Subalpine population (Figure 8B).

**Figure 8.**
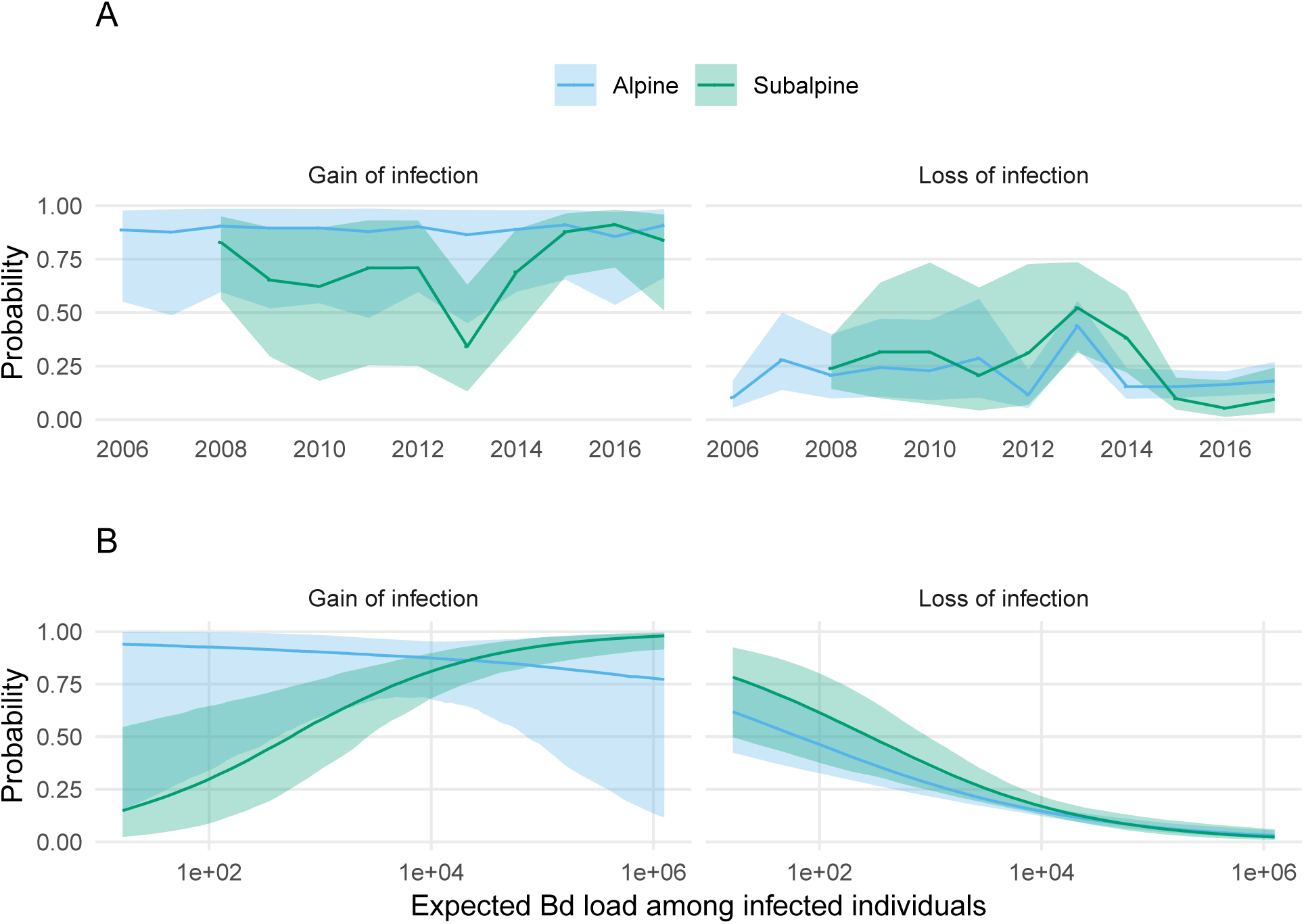
A. Estimated transition probabilities between the uninfected and infected adult classes at both sites in each year. Ribbons represent the 90% posterior credible interval, with the midline representing the posterior median. Dots connected by lines represent years. B. Estimated transition probabilities from the uninfected adult class to the infected adult class and vice versa as a function of expected Bd load among infected adults. The line represents the posterior median, and the ribbons indicate the 95% credible interval, with colors differentiating study sites.

## Discussion

The modeling approach developed here provides a quantitative means to assess how climate and disease affect demographic rates and population dynamics in hard-to-sample host populations. This builds on previous research that incorporated individual-level covariates in mark-recapture models (Pledger et al. 2003, Royle 2008, Gimenez and Choquet 2010, Ford et al. 2012), and that evaluated effects of Bd load at the individual level in wild populations (Spitzen-van der Sluijs et al. 2017). Our approach is unique in part because we treat unobserved infection status and intensity as parameters, rather than using imputation to backfill missing intensity values. Accounting for uncertainty in individual-level infection intensity in a Bayesian framework simplifies uncertainty propagation to key parameters including detection and survival probabilities. This may provide a better understanding of how individual traits relate to demographic processes and population dynamics, especially when effects of disease on detectability lead to biased capture histories.

Overall, results indicated that the introduction of frogs to Alpine was successful in establishing a large self-sustaining population, but unsuccessful at Subalpine. The success of the introduction effort at Alpine seems to be driven by relatively low Bd loads, high adult survival, and large recruitment pulses into the adult population following mild winters. In contrast, the failure of introductions into Subalpine was associated with higher Bd loads, lower adult survival, and smaller, infrequent recruitment pulses. The estimated effect of Bd load on the survival of infected adults was similar in both populations, but loads on average were higher at Subalpine. Given the negative effect of Bd load on frog survival, the cause of higher loads on frogs at Subalpine is worthy of additional study. In addition, the fact that the effect of Bd load on survival of infected adults was similar in both populations, but loads on average were higher at Subalpine also suggests that the observed introduction outcomes cannot be explained by differences between the two populations in frog tolerance of Bd infection within summer seasons (Råberg et al. 2009, Wilber et al. 2017).

The two study lakes were relatively distant from each other and located in catchments with different environmental characteristics, but despite this Bd loads were temporally correlated across the two sites. The synchrony in expected load is worth further investigation, as it may indicate environmental forcing of disease dynamics, e.g., variation due to temperature (Phillott et al. 2013, Cohen et al. 2017), but see (Knapp et al. 2011), or some form of connectivity among sites, although the latter possibility seems unlikely. Future studies might also seek to better disentangle landscape-wide factors driving synchrony and local factors that drive differences in mean load, particularly in the context of known mechanisms of disease-induced extinction such as non-density dependent transmission (Rachowicz and Briggs 2007, Orlofske et al. 2018), nonamphibian reservoir hosts (McMahon et al. 2013), and small equilibrium densities in the presence of disease (De Castro and Bolker 2005).

Although between-population differences in Bd load provide a straightforward explanation for the lower survival of infected adults at Subalpine compared to Alpine (Briggs et al. 2010), it is less obvious why uninfected adults also had lower survival at Subalpine. Low uninfected survival may result from differences in habitat characteristics and/or the abundance of terrestrial or aquatic predators that prey on adult frogs. For example, in 2013 ten translocated frogs at each site were tracked for several weeks using radio telemetry. At Subalpine, two of these frogs were preyed on by garter snakes (*Thamnophis elegans*, (Jennings et al. 1992)) and no evidence of predation was observed at Alpine.

We found that infected adults were easier to detect than uninfected adults, and at Alpine, infection load increased detection probability among infected individuals. This could relate to infection-related behavior that makes frogs easier to find and/or capture (Johnson 2002). For example, some amphibians increase feeding activity when infected with Bd (Hess et al. 2015). If this is the case for *R. sierrae*, we might expect frogs to spend more time around the lake edge, where they would be more conspicuous to observers. Some previous studies have assumed equal detectability among infected and uninfected frogs (Briggs et al. 2010, Stegen et al. 2017), while others have not (Retallick et al. 2004, Murray et al. 2009, Phillott et al. 2013). Our results indicate that even accounting for unequal detectability between uninfected and infected animals may not be sufficient, because Bd load may further affect detectability among infected individuals. Future studies should evaluate conditions under which disease impacts on detectability could bias estimates of demographic rates in wild populations.

The recruitment model developed here may be somewhat simplistic in its focus on adults. In reality, recruitment into the adult population is a function of dynamics in the subadult and larval populations. In particular, in mountain yellow-legged frogs and many other frog species Bd infection imposes heavy mortality during metamorphosis, but disease effects on larval stages are relatively weak (Rachowicz and Vredenburg 2004, Rachowicz et al. 2006). Therefore, to fully understand how factors such as winter severity and disease influence population dynamics by way of affecting larvae, subadults, and adults, future efforts might focus on incorporating elements of integral projection models (Wilber et al. 2016) and integrated population models (Schaub and Abadi 2011) with hidden Markov models similar to those we developed in this study. A joint model for larval, subadult, and adult population dynamics would be better able to account for time lags resulting from the 1-4 year larval development period of this species (Vredenburg et al. 2005). For example, one severe winter could have long-lasting effects if it slows the development of larvae so that they develop in three rather than two years, but this effect might not be detectable as a signal for adult recruitment until four years after the severe winter, depending on the growth rate of subadult frogs.

Our results provide important insights into causes of population establishment or likely failure for the endangered *R. sierrae*. The two study populations showed very different patterns of adult survival and recruitment, and in particular, within a population the estimated survival of introduced cohorts was remarkably consistent across all introductions. Although the generality of results obtained from these two populations need to be assessed using data from mark-recapture efforts at other reintroduced populations, the results suggest that the estimated survival of reintroduced frogs could provide an early indication of the site-specific probability of introduction success. In addition, in the future, when survival estimates of reintroduced frogs are available from a larger number of populations, these estimates could eventually allow the identification of site characteristics associated with likely introduction success or failure. This predictive ability would greatly increase the effectiveness of mountain yellow-legged frog recovery efforts.

## Acknowledgments

We thank the following people and institutions for important contributions to this study: numerous people who assisted with the field work (especially N. Kauffman, A. Killion, A. Lindauer, J. Maurer, and S. Ostoja); R. Chen, K. Rose, and M. Toothman for qPCR assistance; C. Briggs and M. Wilber for reviewing an earlier draft of the manuscript; and Yosemite National Park (R. Grasso, H. McKenny, and S. Thompson), U.S. Geological Survey - Western Ecological Research Center (M. Brooks and S. Ostoja), and the University of California Natural Reserve System - Sierra Nevada Aquatic Research Laboratory (DOI: 10.21973/N3966F) for logistical support and valuable discussions. Research permits were provided by Yosemite National Park, U.S. Fish and Wildlife Service, and the Institutional Animal Care and Use Committee at the University of California, Santa Barbara. This project was funded by grants from the National Park Service (to R.A.K), Yosemite Conservancy (to Yosemite National Park and R.A.K.), U.S. Geological Survey (Ecosystems Mission Area - Natural Resource Preservation Program, to M. Brooks and R. Knapp for the project titled “Factors influencing reintroduction success of the endangered mountain yellow-legged frog”, and National Science Foundation (DEB-1557190, to C. Briggs and R. Knapp). We also thank H. Ito for translating mark-recapture model specifications from JAGS to Stan, as this greatly simplified our implementation.

## Author contributions

R.A.K. designed and performed the research; R.A.K. assembled and maintained the study data set; M.B.J. and R.A.K. developed the models, M.B.J. analyzed the data; M.B.J. and R.A.K. wrote the paper.

## Appendices

### Appendix A: Joint distribution factorization

For completeness, we specify the factorization of the joint distribution of data and parameters (the unnormalized posterior density) below. We represent all unknowns specific to the detection model as a vector *θ*^(*p*)^ = (*α*^(*p*)^, *β*^(*p,x*)^, *β*^(*p,*+)^, *β*^(*p,z*)^), parameters specific to the probability of entry model as a vector *θ*^(λ)^, to the probability of recruiting into the infected adult class given that an individual has entered the population as *θ*^(*γ*)^, to uninfected survival as *θ*(*φ*^-^), and to the infected survival model component as *θ*^(*ø*+^). Concatenating these two vectors gives a vector that contains all unique survival model parameters: *θ*^(*φ*)^ = (*θ*(*φ*^-^), *θ*^(*ø*+^)). Last, we represent parameters unique to transitions from the infected to uninfected class as *θ*(*η*^-^), and from the uninfected to infected class as *θ*^(*η*+^), with both concatenated as *θ*^(*η*)^ = (*θ*(*η*^-^), *θ*^(*η*+^)). The resulting joint distribution is:

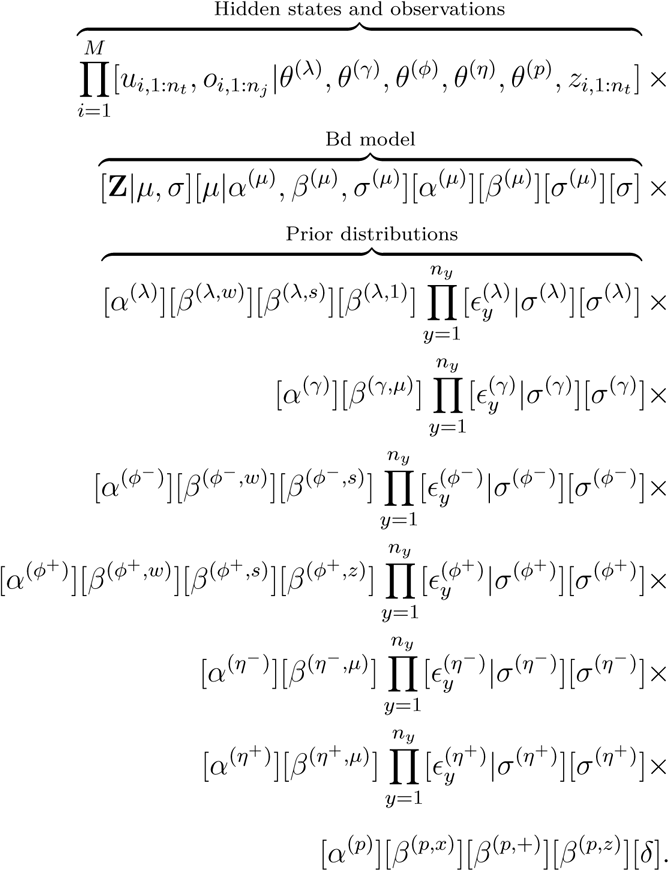

### Appendix B: Forward algorithm description

Parameter estimation for this model is made somewhat difficult by the presence of discrete parameters (hidden states). We address this issue by using the forward algorithm, which does not require sampling from the discrete state space, to compute the joint probability of hidden states and observations (Zucchini et al. 2016). To describe this algorithm, we first consider the case of one individual. We would like to compute [*u*_*i,*1:*nt*_, *o*_*i,*1:*nj*_ *|…*] suppressing dependence on detection and transition parameters for compactness) for the individual with state and capture history shown in Figure 1. We can factor this joint probability as follows:

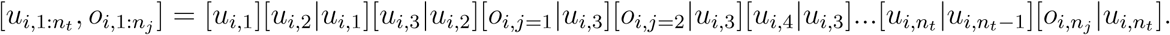

If all of the unknown states were known, this would be as simple as extracting the relevant probabilities from Ψ_*i,t*_ and Ω_*i,j*_. Assuming that all individuals in the first primary period are in the “not recruited” class (*u*_*i,*1_ = 1 *∀i*) implies that [*u*_*i,*1_] = (1 0 0 0), where each element in the row vector represents the probability of being in hidden state 1, 2, 3, and 4, respectively. If we define: **P**(*o*_*i,j*_) = diag(Ω_.,*i,j,o*_) to be the square matrix acquired by placing the elements of column *o*_*i,j*_ from Ω_*i,j*_ along the diagonal (with zeros elsewhere), the forward algorithm provides the joint distribution of hidden states and observations as follows:

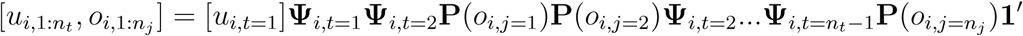

where **1´** is a column vector of ones. More generally, we can compute this probability by defining **B**_*i,t*_ = Ψ_*i,t-*1_ II*j∼t* **P**(*o*_*i,j*_), where *j* ∼ *t* indicates surveys that took place in primary period *t* (if no surveys took place, then **B**_*i,t*_ = Ψ_*i,t-*1_). Then bringing back dependence on all other detection and transition parameters into our notation, we can compute the joint probability of hidden states and the observation history compactly as:

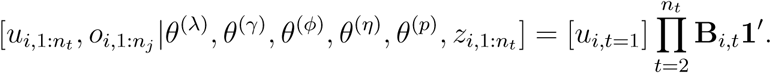

